# Assessing fish diversity in the Amazon: The impact of primers and reference libraries in eDNA surveys

**DOI:** 10.1101/2025.11.07.686145

**Authors:** Shizuka Hashimoto, Sarah V. Carvalho, Sonny de S. C. Miranda, Valéria N. Machado, Lucélia N. Carvalho, Cláudia P. de Deus, Izeni P. Farias, Tomas Hrbek

## Abstract

Selecting primers for DNA amplification poses a challenge in environmental DNA (eDNA) studies, particularly in biodiverse tropical ecosystems. Researchers must balance taxonomic coverage, primer specificity, and study feasibility when choosing between a single primer pair and multiple primer pairs. Empirical tests comparing these strategies are essential for improving eDNA protocols, enhancing the detection of aquatic biodiversity, and supporting conservation and environmental monitoring efforts. This study aimed to evaluate how different approaches affect the specificity and efficiency of detecting fish species in eDNA samples from the Teles Pires River basin. We (1) synthesized modified versions of primers commonly used in ichthyofaunal surveys; (2) compared the results generated by these modified primers with the standard primer versions; and (3) evaluated the species detection using two reference sequence databases: a global taxonomic database (GenBank) and a locally constructed taxonomic database containing samples of fish species previously collected in situ. The 12S-V5_mod primers detected the most fish species across the databases (67, 104, and 130 in the Midori2, Local, and joint databases, respectively), followed by the MiFish_mod primers, which detected 47, 101, and 114 species. Finally, the MiFish primers detected 39, 73, and 78 species. The 12S-V5_mod primers also had lower taxonomic specificity of fish species and amplified a broader range of vertebrate taxa than the MiFish_mod primers. The modified primers outperformed the MiFish primers, and detectability nearly doubled when the Local reference database was used. These results speak to the critical need for constructing reference databases of regional biodiversity, and the need to incorporate multiple primer pairs in eDNA studies.

## Introduction

The Amazon basin is home to the world’s most diverse ichthyofauna, with estimates exceeding 5,000 species [1]. This estimate is growing rapidly. Of the 407 new species of fishes described in 2004, 260 were freshwater and 84 from South America, and most of them from the Amazon basin [2]. Of the 5,000+ species of Amazonian fishes, only 254 have DNA reference sequences for the Miya 12S region, and 324 have DNA reference sequences for the Riaz 12S region [3] in GenBank Release 259. The Miya 12S region taxa are a perfect subset of the Riaz 12S region taxa. The 324 species with 12S DNA reference sequences represent only 5% of the described fish species diversity of the Amazon basin, presenting a significant impediment in our ability to conduct biodiversity surveys and monitoring based on environmental DNA (eDNA) analyses. This is especially worrying as this rich aquatic biodiversity is increasingly threatened by human activities, including the intensive use of water resources for agricultural expansion, urbanization, and the construction of hydroelectric dams, all of which contribute to significant biodiversity loss [4–6].

The expansion of agriculture and deforestation has led to increased river silting, habitat degradation, and water contamination from pesticides, all of which directly impact fish populations [4]. For example, the Teles Pires River basin is located in a region characterized by intense agricultural activity [7,8], significant sport fishing [9], and several hydroelectric projects [10]. These activities have direct and negative effects on the local aquatic fauna.

The rapid transformation of aquatic environments due to anthropogenic pressures coupled with biodiversity knowledge gaps demands the adoption of more efficient and faster monitoring methods for monitoring and identifying fauna. Traditional methods, while essential, often struggle to keep pace with the speed of biodiversity loss, as they rely on physical sampling and detailed taxonomic identification by qualified experts, which is time-consuming and costly. In this context, the analysis of environmental DNA (eDNA) emerges as an innovative and highly sensitive approach for monitoring aquatic biodiversity [11]. Environmental DNA analyses enable the detection of species presence from traces of genetic material released into the environment, such as cell fragments, excrement, mucus, and decomposing matter [12], making identification faster, less invasive, and potentially more comprehensive than conventional methods. Therefore, eDNA represents a promising tool for the early detection of invasive species, monitoring threatened populations, and assessing the health of aquatic ecosystems [13], thereby aiding biodiversity conservation and management plans.

The analysis of eDNA from water has emerged as a powerful tool for conducting more detailed and efficient assessments of fish communities [14,15]. Additionally, eDNA allows for the analysis of the temporal dynamics of fish communities, including seasonal and long-term variations [16], and provides data on the relative abundance of different species [17]. This methodological advancement represents a significant step toward understanding aquatic biodiversity and enhancing conservation and environmental management strategies.

In eDNA studies, the mitochondrial 12S rDNA gene is commonly used for fish species identification due to the specificity of the primers used and the effectiveness of the amplified region in distinguishing among different taxa [18–20]. While the 12S gene is prevalent, some studies incorporate other gene regions, such as COI, 16S, and 18S [21–24]. The most frequently used primers for sampling aquatic environments and fish include MiFish-U [25] and 12S-V5 [26], each amplifying different regions within the 12S gene.

The choice of primers for DNA amplification poses a challenge in eDNA studies, particularly in diverse ecosystems like the Neotropics which are taxonomically very diverse. For example the 3’ ends of the universal primers developed by Miya et al. (2015) for fishes have mismatches with characiform 12S rDNA. This will result in amplification biases that favor non-characiform lineages and will lead to an underestimation of actual biodiversity [27]. Specifically, while Characiformes represent about 43% of the Amazon basin’s total fish species diversity, in the eDNA survey of the lower Javari River basin, characiform fishes represented only 27.1% of the detected OTUs [28]. Some of these biases can be mitigated by the use of multiple primer sets which can enhance taxonomic coverage and improve the detection of species with varying target sequences [18], however, at the cost of additional time, laboratory and analytical effort and expense. Solutions can also be sought in redesigning primers to better match and amplify local biodiversity.

The success of any eDNA study is predicated on the availability of a locality informative reference database, a recurring issue in studies comparing primers where primer “success” is predicated on the reference database used [29–31]. With only about 5% of all Amazonian basin fishes having one or both of the commonly used 12S regions included in public databases, biodiversity monitoring via eDNA will always be lacking until more effort is put into generating curated reference databases. In an ichthyofaunal survey of the lower Javari River in the western Amazon basin, Santana et al. [28] sampled 443 species by traditional methods—of which 60 were new to science—and detected 222 operational taxonomic units of which 58 could be assigned to species using a 98.5% similarity criterion and the publicly available MiFish DB ver. 36. However, only 17 species were detected by both physical and eDNA surveys. The development of robust reference databases that incorporate sequences from locally collected species has significantly improved the accuracy of species detection compared to public databases [32].

In this study, we synthesized two primer cocktails which also capture the taxonomic diversity of Amazonian fishes to evaluate and compare their performance with the MiFish primer, which is widely used in eDNA studies. Our goal was to assess the specificity of the sequences (% of taxa assigned to species level in Fig. 3), informativeness (number of species detected in Fig. 3) and efficiency (% on target sequences in Table 2) in detecting vertebrates in general and fish specifically in the Teles Pires River basin. Specifically, we aimed to: a) compare the original primer from Miya et al. [25] with our modified primers for Amazonian fishes; b) evaluate the efficiency of these primer sets using both the global taxonomic database, GenBank, and a locally constructed taxonomic database to determine how much GenBank can detect and how much the local reference enhances the accuracy of detecting local diversity; and c) make publically available a new set of primers which will make the study of Amazonian biodiversity more efficient and comprehensive.

## Materials and methods

### eDNA study area and sampling

The confluence of the Teles Pires and the Juruena rivers forms the Tapajós River, a major clear-water tributary on the right bank of the Amazon River [33]. The Teles Pires River spans a total length of 1,457 km, with the majority of its drainage area located in the state of Mato Grosso (1,370 km) and a smaller portion in the state of Pará. The Teles Pires River is characterized as a clear-water river due to the typical rock formations of the Brazilian Shield, which drain waters with low amounts of sediment and organic matter, resulting in low electrical conductivity. The hydrological cycle exhibits a flood pulse pattern, with the rainy season occurring from November to February and the dry season from May to October, reflecting the region’s seasonality [34].

For this study, we sampled at 22 localities along the Teles Pires River (Fig 1) during the months of July and August, 2021. At each locality we collected three replicates using a Van Dorn bottle and stored them in labeled 500 mL sterile bottles, kept away from light and heat. We then filtered each sample using a 50 mL disposable syringe fitted with a 0.45 μm Sterivex-GPTM PES filter (Merck Millipore, Inc., Darmstadt, Germany) attached to the tip. We completed all sample filtration within 24 hours of collection to prevent degradation of the DNA present in the samples. After each filtration, the equipment were sterilized with 10% sodium hypochlorite for 10 minutes and rinsed with distilled water, following the protocol of Handley et al. [16] to avoid the risk of contamination. The filters were stored in an ATL buffer (Qiagen), sealed, labeled, and stored away from light and heat until DNA extraction.

**Fig 1.**
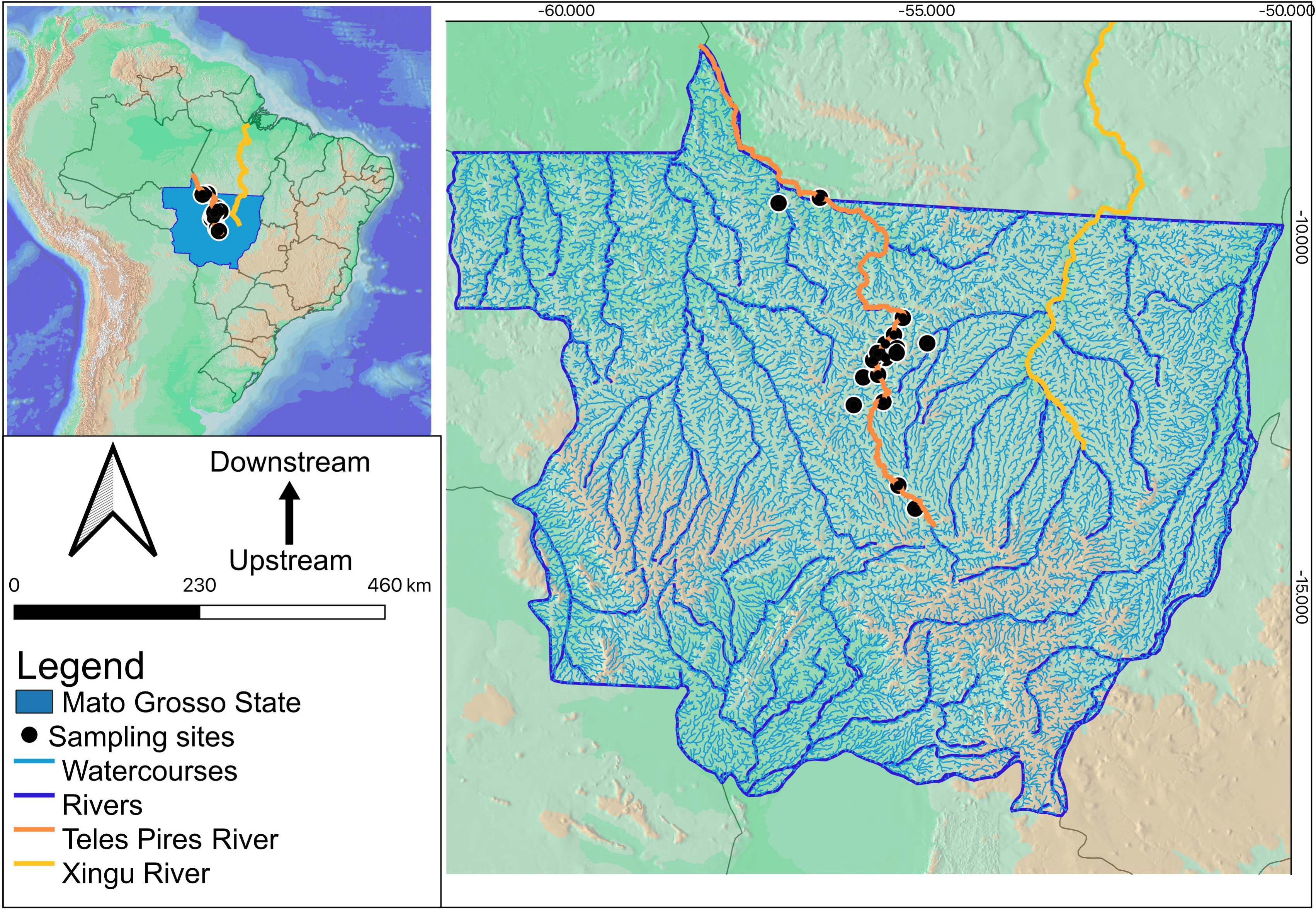
Sampling sites along the Teles Pires River in Brazil.

**Fig 2.**
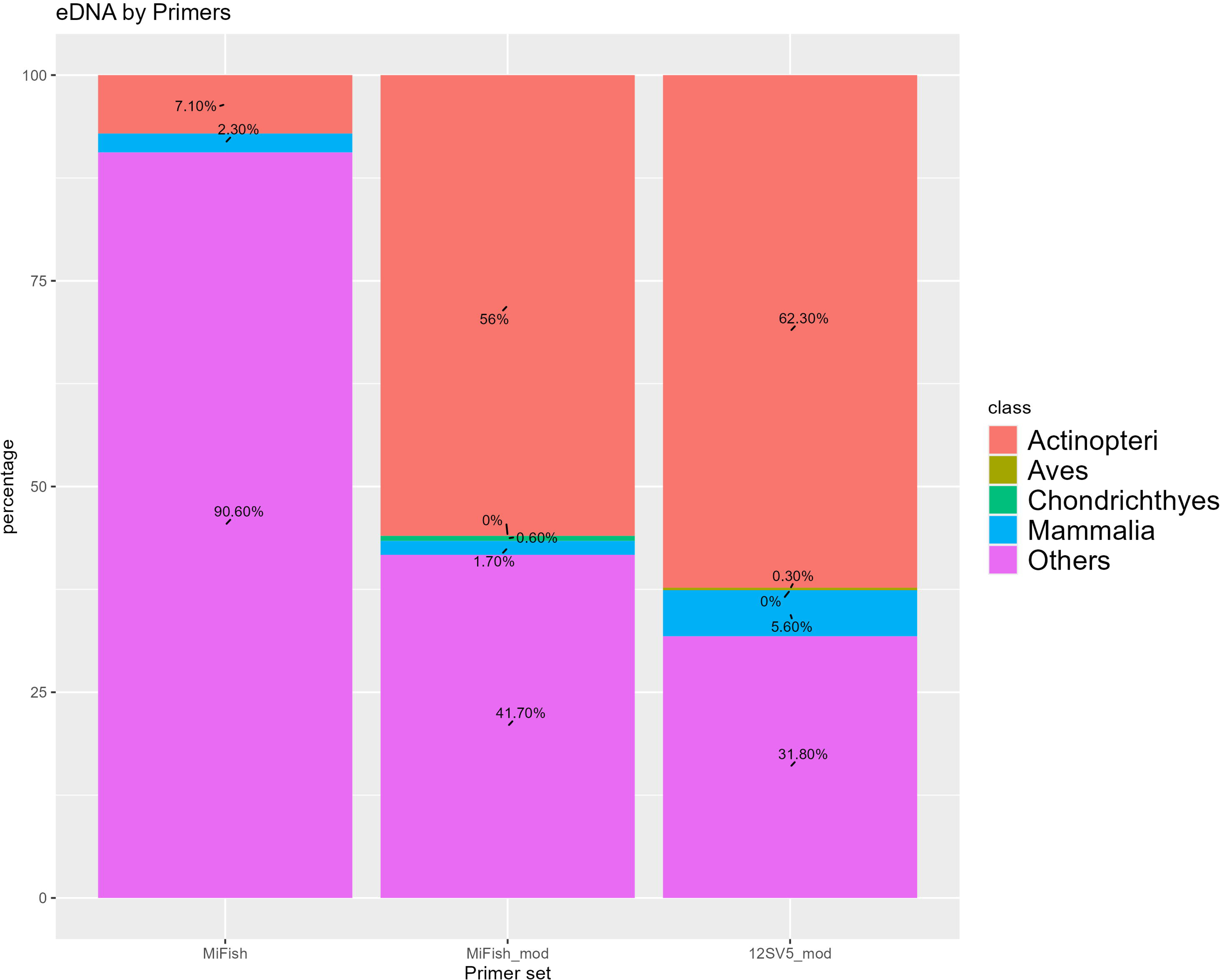
Proportion of reads assigned to vertebrate class levels for the three metabarcoding primer sets.

### Laboratory processing

We extracted DNA in a dedicated eDNA extraction room, separate from the PCR room. Each extraction began with cleaning the room using 70% alcohol, followed by bleach for decontamination. We also disinfected all instruments, including pipettes, scalpels, and tweezers used in the DNA extraction steps with alcohol and bleach, and exposed them to UV light for 30 minutes as a final cleaning step.

We then extracted DNA from the filters using the DNeasy PowerSoil Kit (Qiagen), with a modification in the digestion step, which involved digestion at 60 °C for 2 hours on a shaker, followed by quantification using a NanoDrop 2000 spectrophotometer (Thermo Scientific).

We amplified each gene fragment using a two-step PCR protocol. The first step targeted the amplification of a mitochondrial 12S rDNA gene fragment, while the second step involved ligating sequencing adapters for sample identification during high-throughput sequencing. We carried out the first PCR in a final volume of 25 µl, which included: 17.45 µl of ultrapure water, 2.5 µl of 10X buffer (containing 500 mM Tris-Cl pH 9.2, 160 mM ammonium sulfate, 0.5% Brij 58, and 35 mM magnesium chloride), 2.5 µl of dNTPs (at a concentration of 2 mM), 0.75 µl of each 2µM primer (forward and reverse), 0.05 µl of OmniTaq polymerase (DNA Polymerase Technology, Inc.), and 1.0 µl of environmental DNA (eDNA). In addition to the original MiFish primers [25], we also amplified the eDNA using modified versions of the MiFish and 12SV5 primers [26] (Table 1). These modifications were based on observed sequence variation in the 12S region of Amazonian fishes deposited in GenBank.

**Table 1.**
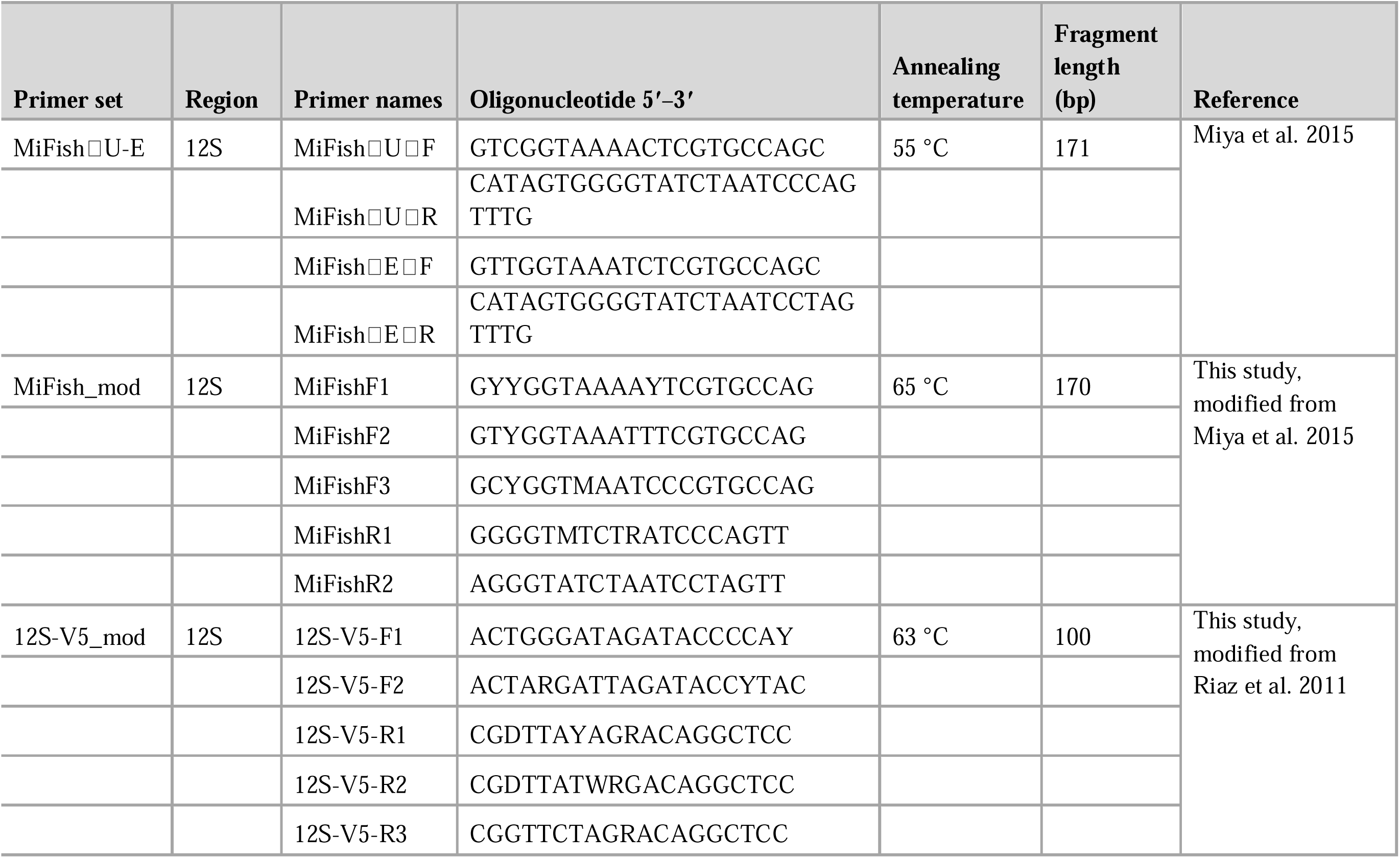
Primer set used in this study without NGS Illumina adapters.

The PCR cycling conditions were as follows: initial denaturation at 94 °C for 1 minute; followed by 40 cycles of denaturation at 94 °C for 5 seconds, primer-specific annealing at a designated temperature (Table 1) for 30 seconds, and extension at 68 °C for 10 seconds. We performed each PCR reaction, including negative controls, in triplicates to minimize amplification biases caused by stochastic variation in early PCR cycles. We visualized each PCR product on a 1% agarose gel stained with GelRed (Biotium), and then pooled the triplicate reactions into a single composite sample and bead purified it [35] for subsequent indexing PCR.

We used a dual indexing strategy to individualize each sample. In the indexing PCR we used the same conditions as in the first PCR except we used one Q5 and one Q7 primer at 55C annealing temperature, 3 µl of the first PCR product, and 15 PCR cycles. We evaluated the PCR reactions on a 1% agarose gel, and then purified them using BOMB paramagnetic beads [35]. After quantifying each purified PCR product using the Qubit fluorometer, we pooled all PCR products equimolarly for sequencing. We sequenced the pooled library on the Illumina HiSeq 4000 platform (Illumina, Azenta Life Sciences, USA).

### Construction of the local reference library

We constructed the Local DNA reference library using samples of fish specimens collected between 2016 and 2019 as part of the ichthyofauna monitoring program associated with the Sinop Hydroelectric Power Plant (HPP). All specimens were identified to the species level, cataloged, and deposited as voucher specimens in the Ichthyological Collection of the Southern Amazon Biological Collection, housed at the Tropical Ichthyology Laboratory (LIT-ABAM) in Sinop, Mato Grosso, by the ichthyological staff of LIT-ABAM [36]. Tissue samples of these specimens were deposited in LIT-ABAM, CTGA tissue collection of UFAM or the genetic resources collection of INPA. Of the 354 species of fishes recorded by Carvalho et al. [36] for the middle and upper Teles Pires River, 173 species had tissues available and were included in the Local DNA reference library, representing 49% of the species collected in the faunal surveys prior to the construction of the Sinop Hydroelectric Power Plant [36].

To complement the Local database, we also included additional species from the Xingu River basin, some of which might be shared between the two basins—documented in the studies by Carvalho et al. [37] and Ximenes et al. [38]. Muscle tissue samples from these additional specimens are archived in the Animal Genetics Tissue Collection of the Laboratory of Animal Evolution and Genetics at the Federal University of Amazonas (CTGA/LEGAL/UFAM). On average, two individuals per species were included in the local database, resulting in a total of 390 species, representing 171 genera, 47 families, and 12 orders (Table S1).

The collections had the Special Fishing License (LEP 352/2016 and LEP 855/2018) from the State Secretariat for the Environment – SEMA. This research was carried out in accordance with the rules and regulations of Ministério do Meio Ambiente, Instituto Chico Mendes de Conservação da Biodiversidade under permit number SISBIO 76359-1. Access to the genetic heritage associated with this study was registered in the National System for the Management of Genetic Heritage and Associated Traditional Knowledge (Sistema Nacional de Gestão do Patrimônio Genético e do Conhecimento Tradicional Associado – SISGEN) under accession number AB8314D.

We used the Midori2 raw fasta GB259 database [39] as our global reference library. We standardized the taxonomy of both the Local as well as the Midori2 reference database to a seven level taxonomy using TaxonKit [40] and validated the taxonomy of fish taxa using the Eschmeyer’s Catalog of Fishes [41]. Finally, we simplified the Midori2 database fasta headers to Genbank IDs only, and using this simplified Midori2 fasta, the Local database fasta and the merged fasta, we generated three local BLAST+ databases.

### Bioinformatics processing

The raw data were processed using the package barcode_splitter v0.18.6 [42], which sorted the samples into folders based on their index sequences. From the raw reads, we extracted only those sequences which had both the forward and reverse primers on both ends, and then removed these primers with cutadapt v5.0 [43]. The paired-end reads were merged into consensus sequences using PEAR v0.9.11 (Zhang et al., 2014), chimeric sequences were removed and then the merged reads were dereplicated using VSEARCH v2.21.1 [45] and clustered using SWARM v3.1.4 [46]. We then compared the clustered sequences against the Midori2 global reference library [39], a Local reference library, and a merged global/local library using BLAST+ [47].

### Statistical analyses

We processed the BLAST+ output using a custom R [48] script, assigning sequences to taxa based on similarity thresholds following Beatty et al. [49]: ≥97% for species, ≥95% for genus, ≥90% for family, and ≥80% for order, removing all sequences shorter than 99 bp, and all taxa with less than four reads. We visualized the results using ggplot2 [50] and the VennDiagram package v1.7.3 [51]. To estimate species richness and visualize accumulation curves across primers, we used the iNEXT package v3.0.0 [52], based on species abundance and richness data.

## Results

### Primer set performance

From the 66 samples and nine negative controls, a total of 62,278,389 raw sequences were generated. Among the primers, MiFish produced the highest number of raw sequences (29,278,389), followed by 12S-V5_mod (19,685,401) and MiFish_mod (13,093,553). To enable fair comparison of the three primer pairs, we standardized the read counts to the number of reads in the MiFish_mod primer pair which corresponded to an average of 189,761 reads per sample.

Post-filtering retention (on target) rates were 1.06% for MiFish, 19.65% for MiFish_mod, and 20.46% for 12S-V5_mod. When using the Midori2 database for taxonomic assignment, 4.8% of MiFish on target reads (7,027 reads), 25.9% of MiFish_mod on target reads (666,394), and 41.9% of 12S-V5_mod on target reads (1,121,444) were successfully assigned as fish at any taxonomic level. Using the Local reference database, 6.9% of MiFish on target reads (10,207), 56.5% of MiFish_mod on target reads (1,453,323), and 60.1% of 12S-V5_mod on target reads (1,607,832) were classified as fish at any taxonomic level. Combining both databases, 7.1% of MiFish on target reads (10,456), 56.5% of MiFish_mod on target reads (1,454,749), and 62.4% of 12S-V5_mod on target read (1,667,143) were successfully assigned as fish at any taxonomic level. These results indicate that the modified primers (MiFish_mod and 12S-V5_mod) produced cleaner data with significantly higher taxonomic assignment rates and fewer artifacts (Table 2).

**Table 2.**
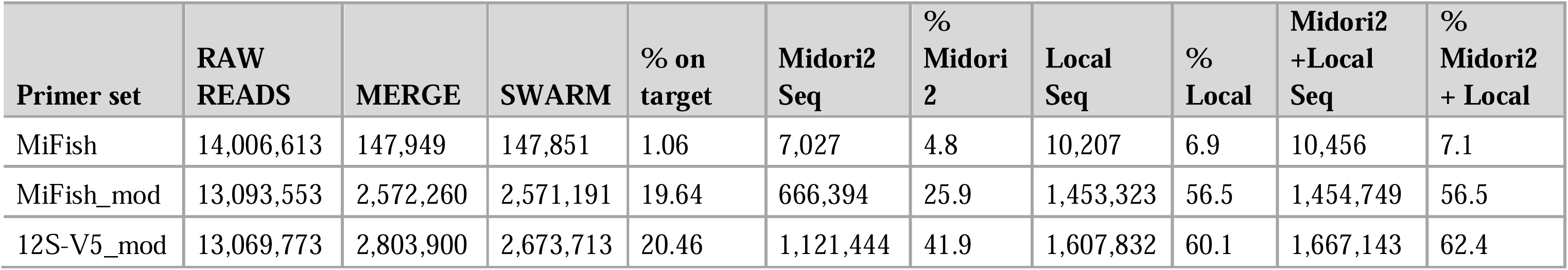
Summary of reads standardization and taxonomic assignments results.

Amplification occurred in the negative controls, generating a total of 242,066 raw reads and 28,824 filtered reads for the MiFish primers; 745,462 raw and 56,933 filtered reads for MiFish_mod; and 543,065 raw and 33,157 filtered reads for 12S-V5_mod. In the negative controls we detected five fish species in the MiFish primers, 16 fish species in MiFish_mod, and 22 fish species in 12S-V5_mod. These sequences were considered potential contaminants in all the samples, thus we removed the equivalent number of contaminating sequences from all samples.

### Primer set x reference database

The MiFish, MiFish_mod and 12S-V5_mod primer sets, in this order, consistently detect an increasing number of taxa across the three reference datasets. The MiFish primer set detects the smallest number of taxa most because it amplifies only a restricted subset of Amazonian fish species, performing especially poorly with Characiform taxa (Fig. 7). When compared with the MiFish_mod primer pair, it detects only 50–60% of the taxa detected by the MiFish_mod primer pair (Fig. 3). At the same time, a greater number of reads can be assigned to species and genus levels, reflecting the intersection of the greater specificity of this primer pair for the taxa that are deposited in public databases. The MiFish_mod primer set amplifies a much broader range of taxa, and this is also reflected in 18% of all taxonomic assignments being made to non-Amazonian taxa when the reference dataset is deficient in the local taxa, such as the Midori2 reference dataset. However, when a local reference database is available, even if it contains only 50% of the expected local diversity, assignments to non Amazonian taxa drops to 4%. The MiFish_mod primer set also has high specificity, with 73.2% taxa being assigned to a species or a genus in the combined Midori2 and Local reference databases (Fig. 3). The 12S-V5_mod primer pair detects the greatest number of taxa but at the costs of lowest specificity with 60.6% of taxa being assigned to a species or a genus in the combined Midori2 and Local reference databases (Fig. 3). The number of detected non-Amazonian fishes was also slightly higher than in the MiFish_mod primer pair (Fig. 3). The number of fish species detected by the 12S-V5_mod and MiFish_mod primer pairs against the Local reference database was effectively the same (104 vs 101 species, respectively, with 53 species being detected by both primers), while against the combined Midori2 and Local reference databases, 129 vs 113 species (53 species shared) were detected using the 12S-V5_mod and MiFish_mod primer pairs, respectively. This is probably explained, at least in part, by the GenBank Release 259—from which the Midori2 reference database was built—having 70 more Amazonian fish species for the Riaz 12S region which the 12S-V5_mod primer pair amplifies than for the Miya 12S region which the MiFish_mod primer pair amplifies. Finally, the 12S-V5_mod primer pair also amplifies more species of other vertebrate taxa than any other primer pair. It amplified species of 21 families of mammals and 22 families of birds, while the MiFish_mod primer pair amplified species of 12 families of mammals and 7 families of birds, and the MiFish primer pair amplified species of 8 families of mammals and no birds (Fig S1). Both the 12S-V5_mod vs MiFish_mod primer pairs clearly perform better than the original MiFish primer pair, however, each performs better in a different way.

**Fig 3.**
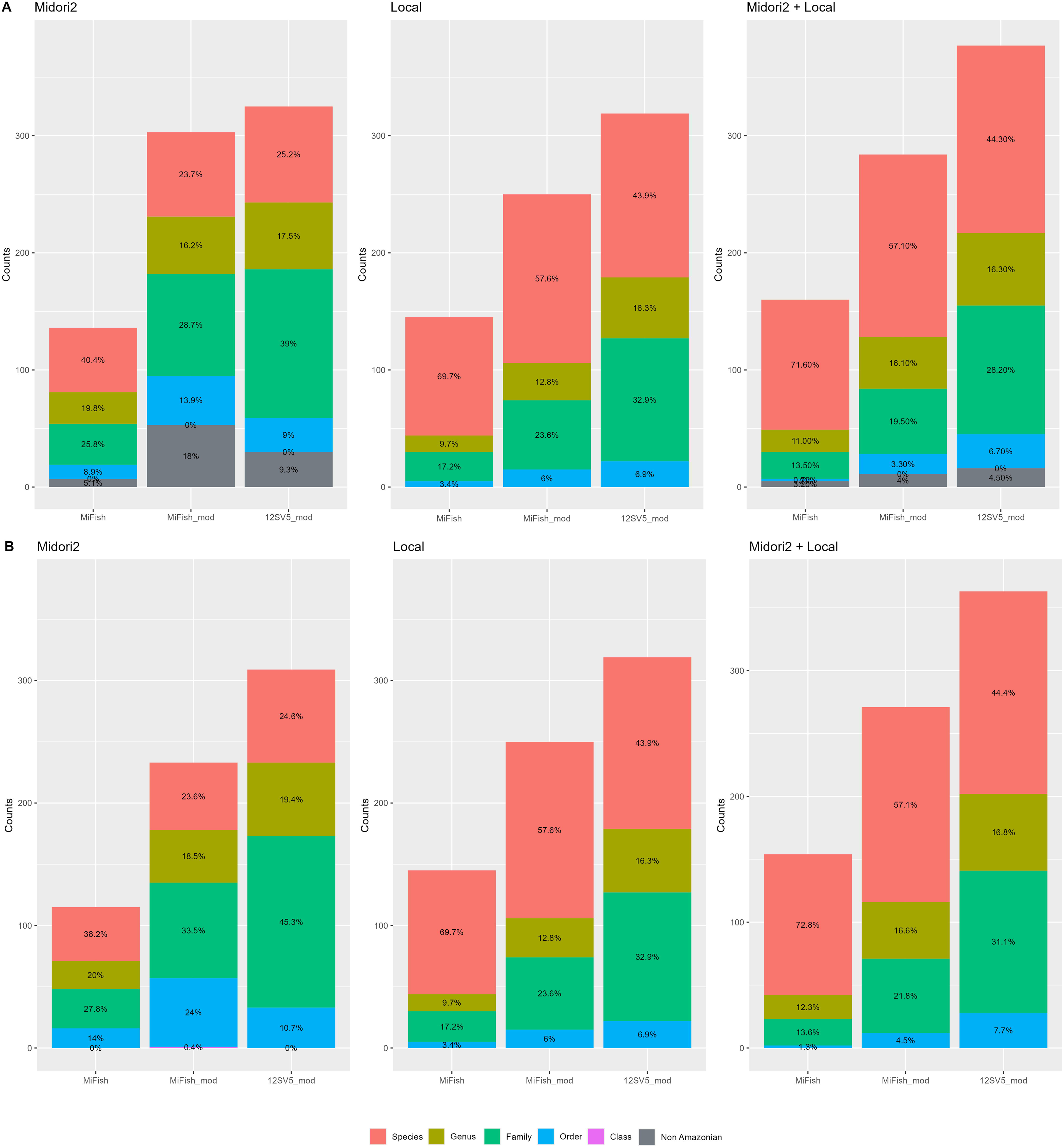
Proportion of reads assigned to each taxonomic level for the three metabarcoding primer sets and across databases. Set A includes all detected taxa at all taxonomic levels. Set B includes only Amazonian taxa.

### Species richness and the collector’s curve

When analyzing species richness, the 12S-V5_mod primers with the merged global/local reference database detected the largest number of species (130 species), and the MiFish primers with the Midori2 database detected the smallest number of species, only 39. The 12S-V5_mod primers were able to detect more species with all three databases: 67, 104 and 130 species corresponding to Midori2, Local and global/local databases, respectively. Next were the MiFish_mods primers with 47, 101 and 114, respectively. And finally, the MiFish primers detected 39, 73 and 78 species, respectively. The modified primers were better than MiFish and species detectability increased when the Midori2 and Local databases were used together, i.e. taxonomically more comprehensive reference databases are better at assigning more sequences to a more specific taxonomic level (Fig 4).

**Fig 4.**
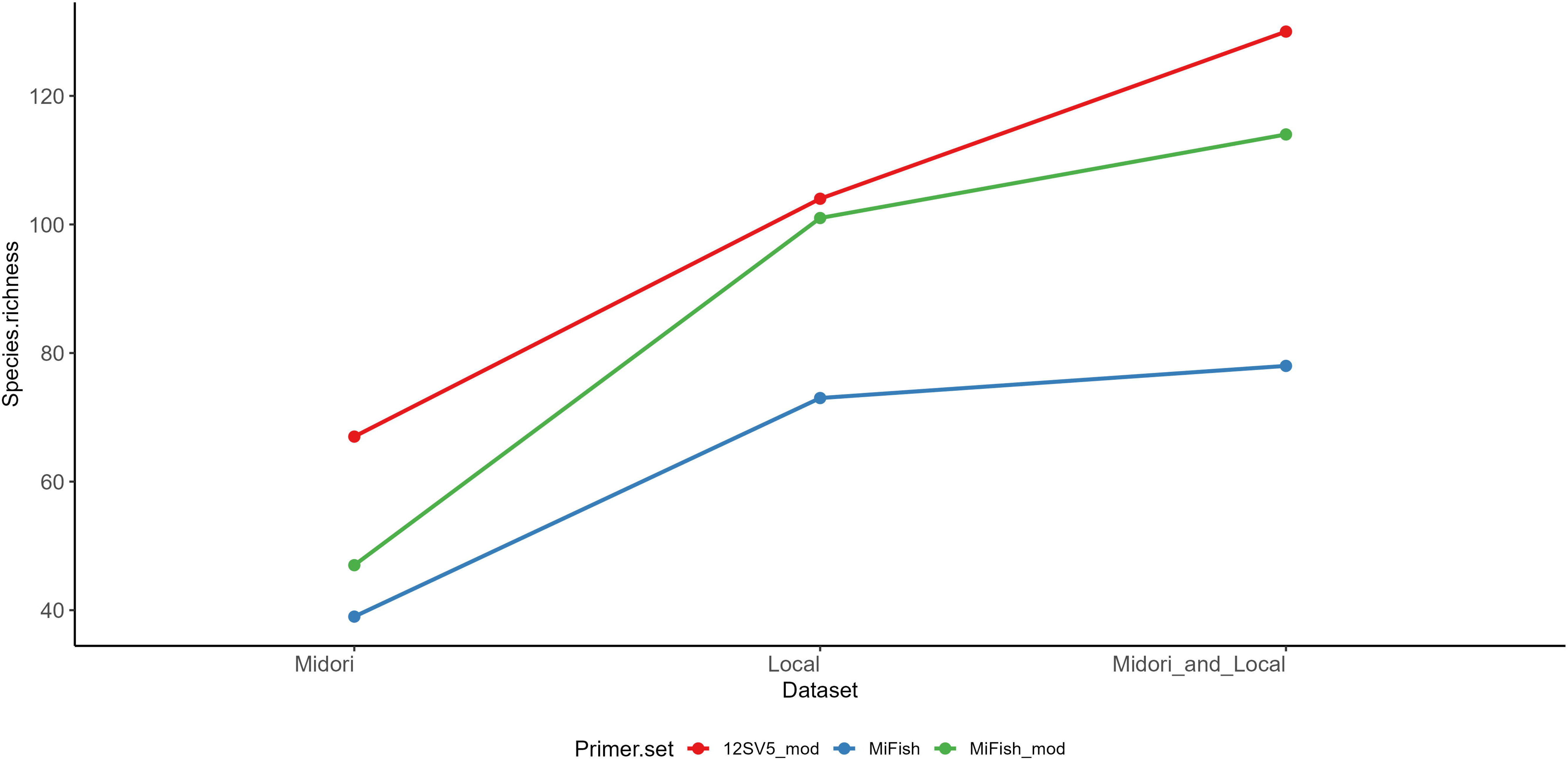
Alpha diversity measures for each primer set and each dataset.

We generated a species accumulation curve to assess the performance of each primer in combination with the reference databases (Fig 5). The results indicate that the 12S-V5_mods primers used with the global and Local reference database were the most effective in detecting species, while the MiFish primers with the Midori2 database performed the worst.

**Fig 5.**
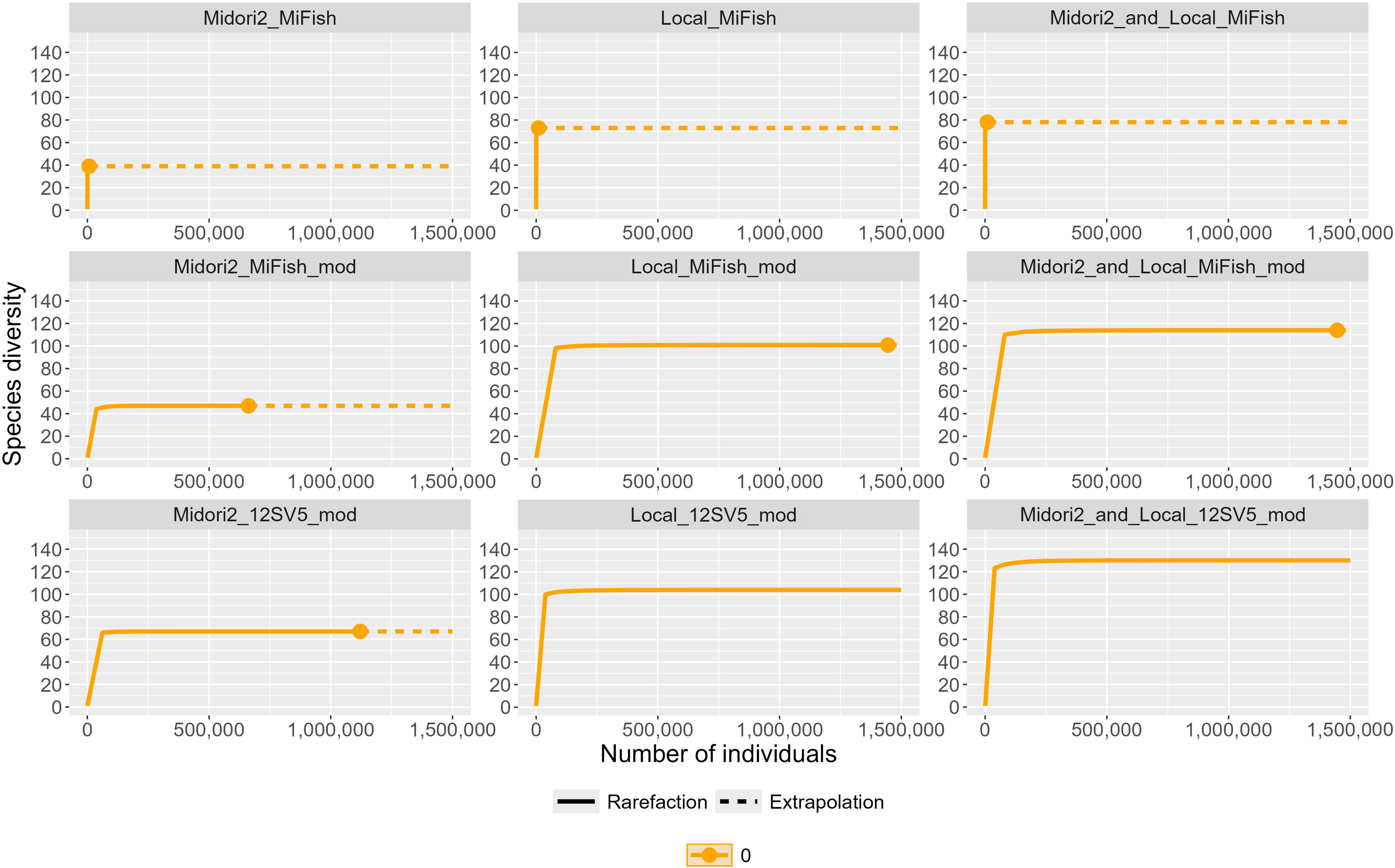
Species accumulation curves for individual metabarcoding primer pairs. Lines represent a sampling number of individuals and dotted lines represent extrapolated curves.

The Local reference database consistently outperformed Midori2, detecting approximately twice as many species. Moreover, combining both databases increased total species detection, as each database contributed unique taxa not found in the other. These findings highlight the critical role of the Local database in enhancing species detection and improving the overall assessment of ichthyofaunal diversity in the upper Teles Pires River basin specifically, and the Amazon basin in general.

### Taxonomic identification

The combined use of all three primer sets and both reference databases (Midori2 and the Local dataset) enabled the identification of 40 families, 118 genera, and 256 species. Of these, 162 species were detected using the Local database, 109 species with the Midori2 database, and only two species—*Hoplias malabaricus* and *Sternopygus macrurus*—were detected in both databases. Within the Local database, the MiFish primer detected 73 species, 10 of which were unique to this primer; 12S-V5_mod detected 104 species, with 46 exclusive detections; and MiFish_mod detected 101 species, 26 of which were unique. A total of 36 species were shared among all three primers. Using the Midori2 database, 39 species were detected with MiFish (13 exclusive), 67 species with 12S-V5_mod (50 exclusive), and 47 species with MiFish_mod (18 exclusive). Considering the combined database, the MiFish primer detected 78 species, including 14 unique to it; 12S-V5_mod detected 129 species, with 71 exclusive detections; and MiFish_mod detected 113 species, 37 of which were unique. These results emphasize the importance of using multiple primers and complementary reference databases to maximize species detection (Fig 6).

**Fig 6.**
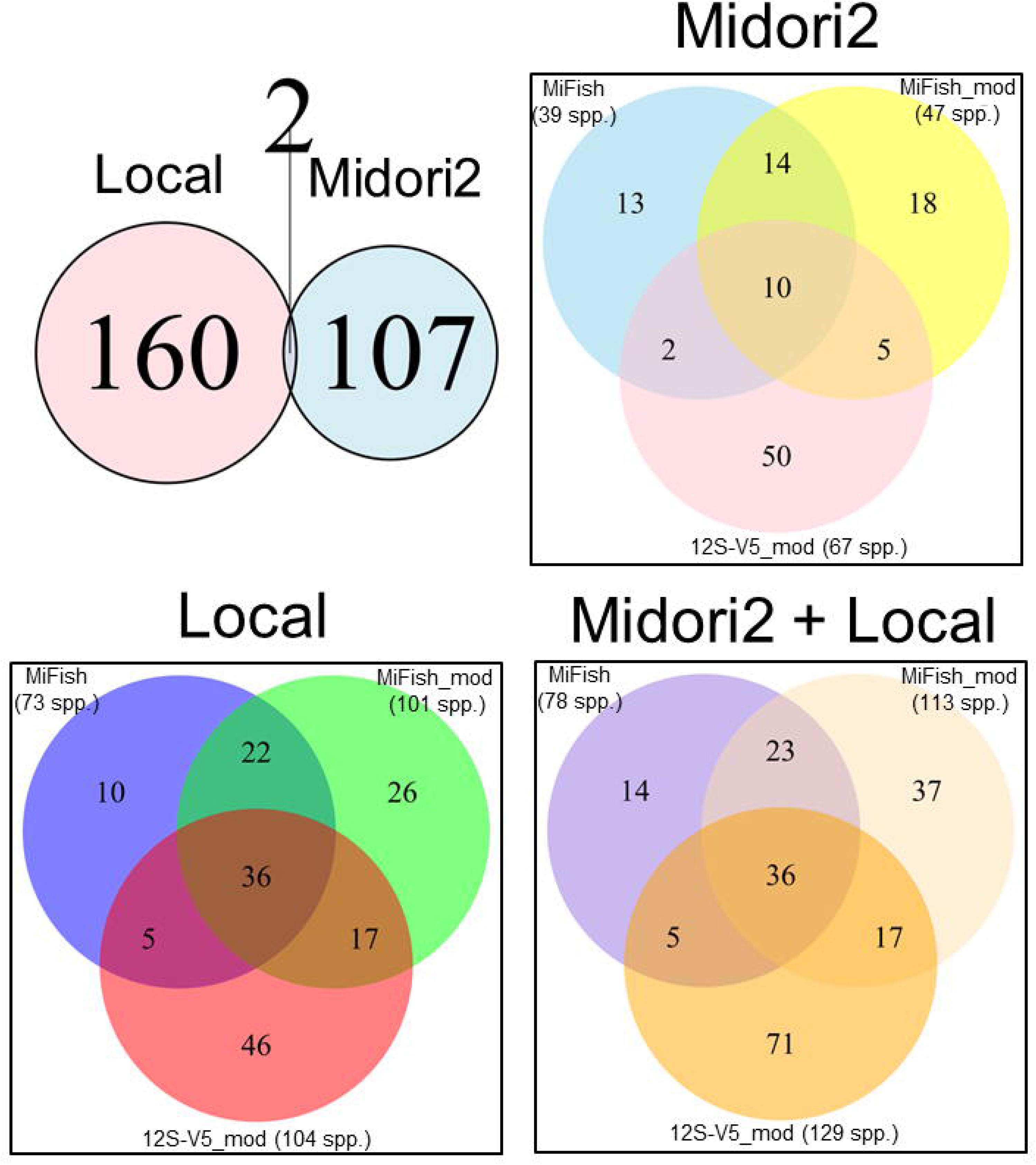
Venn diagrams showing the total species detected in each database, and the contributions of each primer pair (MiFish, MiFish_mod and 12S-V5_mod) on detection of target species.

When comparing taxonomic detection across databases, a total of 25 families were identified using the Midori2 database: 18 families with the MiFish primer pair, 19 with 12S-V5_mod, and 20 with MiFish_mod. In contrast, the Local database identified a broader taxonomic range, detecting 38 families overall—including Potamotrygonidae (Class Chondrichthyes), which was absent from Midori2. Specifically, the MiFish primer pair detected 26 families, 12S-V5_mod detected 31, and MiFish_mod detected 28 families.

When combining both databases, the total number of families detected increased to 40, with MiFish identifying 27 families, 12S-V5_mod identifying 33, and MiFish_mod identifying 29 families.

When compared to the Local and combined databases, the Midori2 database failed to detect any species in any of these 15 families: Acestrorhamphidae, Bryconidae, Chilodontidae, Ctenolucidae, Cynodontidae, Gasteropelecidae, Hypopomidae, Lebiasinidae, Parodontidae, Poecilidae, Potamotrygonidae, Rhamphichthyidae, Rivulidae, Sciaenidae, and Sternopygidae. All but Acestrorhamphidae have representatives in the Midori2 databases, however, these representatives appear to be too distantly related to the species occurring in the Teles Pires River to be detected. In contrast, the Local database did not detect, at the species level, only two families: Apteronotidae (because the Local database does not contain any Apteronotidae) and Synbranchidae (because the local database only contains a sequence of a *Synbranchus royal* from the middle Xingu River which showed 93% similarity to the eDNA sample).

Overall, the 12S-V5_mod primer demonstrated the highest family-level detection, while MiFish showed the lowest. These results underscore the importance of using a comprehensive reference database, particularly a local database, to enhance taxonomic resolution at the family level (Fig 7).

**Fig 7.**
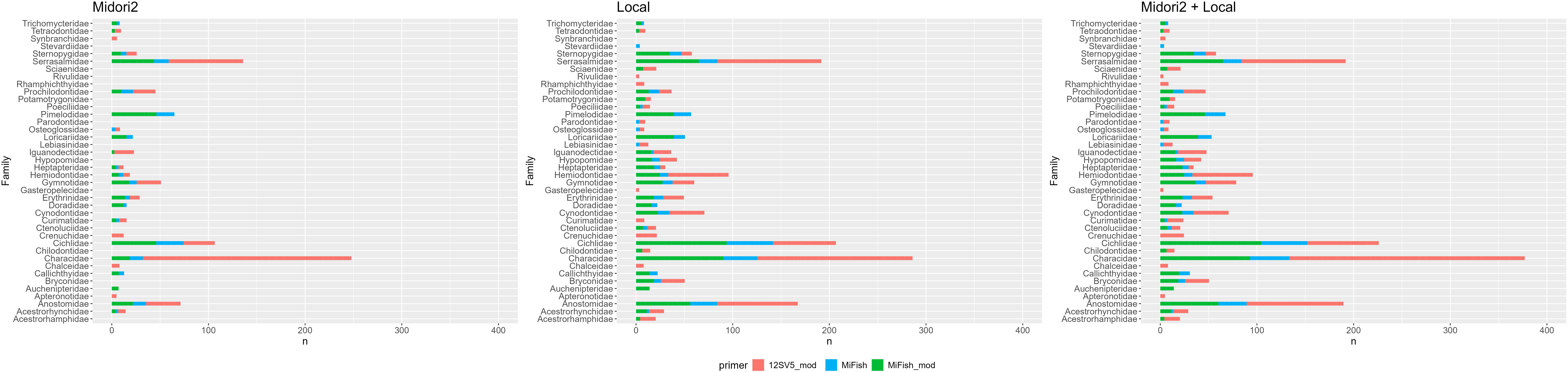
Stacked bar plots showing the relative read abundance o f each primer pair in each family. (A) using Midori2 database. (B) using a Local reference database. (C) using databases together.

It is important to note that the Midori2 database includes a wide range of Eukaryotic taxa, not just the target group (Actinopterygii). As a result, this database also yielded detections of non-target groups, such as birds and mammals. When analyzing the total reads against the complete Midori2 database, we detected a notable number of non-target taxa, including birds from 13 orders, 25 families, and 38 species, and mammals from nine orders, 19 families, and 46 species (see Fig S1 for details). These taxa are primarily detected by the 12S-V5_mod primer set, a modification of the 12S-V5 primer set which amplifies a broad-range of vertebrate taxa [26].

## Discussion

### Primer efficiency

The choice of metabarcoding primers is critical when sampling megadiverse regions such as the Amazon. Although COI reference databases are generally more comprehensive, their application in eDNA studies often results in the detection of relatively few fish sequences and a disproportionately high number of bacterial sequences. This limitation arises from the lack of conserved priming regions. A practical consequence of this is that effective universal primers cannot be designed, and highly degenerate primers generate amplification biases and non-target amplification [18]. In contrast, the 12S gene has proven to be more suitable for eDNA studies, with much higher on-target amplification rates, thereby enhancing the likelihood of detection in environmental samples [19,20,25].

Although the MiFish primers have been used in eDNA studies in the Amazon [28,53,54], our findings indicate that they yield a low percentage of on-target reads (1.06%) compared to the modified primers: MiFish_mod (19.65%) and 12S-V5_mod (20.46%) (Table 2; Fig 2). The MiFish primers also detected only about 50% of taxa that the MiFish_mod and 12S-V5_mod primers detected, although a larger percentage of these taxa were identified to the species level (Fig 3).

The somewhat counterintuitive observation that the MiFish primers showed the greatest taxonomic resolution is explained by their specificity. The MiFish primers being too specific and primarily amplifying taxa and clades which were deposited in public databases or generated by the Miya group—Paleotropical and Paleoarctic fish groups. At the same time, only 1.8 % of all the amplified fragments were on target, and of those only 4.8 % could be assigned to a vertebrate taxon (Table 2). This results in many few vertebrate taxa being detected—only about ½ to ⅗ of the taxa amplified by the MiFish_mod and 12SV5_mod primers, but for those taxa greater proportion are detected at the species level.

When comparing both the MiFish and MiFish_mod primers to the 12S-V5_mod primers, a larger proportion of reads can be assigned to lower taxonomic ranks due to the greater taxonomic informativeness of the MiFish region (170 bp) vs. the 12S-V5 region (100 bp) (Fig 3). However, the total number of species detected by the MiFish_mod and 12S-V5_mod primers is similar (Fig 6).

Although we did not include the original 12S-V5 primer from Riaz et al. [26] in our study, our analyses with the modified 12SV5 primers support the superior performance of the 12S-V5 region over MiFish region amplified with the MiFish primers [25] in detecting species [55–57]. However, our MiFish_mod primers which were modified to better capture Neotropical fish diversity detect approximately the same number of species as the 12S-V5_mod primers (Fig 6). These findings collectively underscore the importance of primer selection in maximizing detection rates and for improving biodiversity assessments in complex tropical ecosystems.

### Importance of a Local database

The comparison between the global reference database—which has the Miya 12S region for 254 species and the Riaz 12S region for 324 species of Amazonian fishes, and the local reference database which has sequences of the Miya 12S and Riaz 12S region for 392 Amazonian fish species, with a focus on the Teles Pires River basin—showed that only two species were shared between them, highlighting the critical role of a local reference library in eDNA-based monitoring of Amazonian ichthyofauna. The Local database permitted the detection of 162 species, compared to the 109 species detected using only the Midori2 database. When both databases were combined, species detection increased substantially to a total of 265 species.

In a previous survey of the ichthyofauna of the Teles Pires River using traditional sampling methods, [33] recorded 11 orders, 42 families, 198 genera, and 355 species of fishes, using a variety of fishing gear such as gillnets, purse seines, longlines, sieves. This survey was carried out over a five year period, four times a year (high, receding, low, rising waters) and at 20 sampling sites distributed along a 220 km long section of the lower portion of the main channel of the Teles Pires River and its tributaries. Similarly, Carvalho et al. [36] registered 354 species in 38 families and nine orders in the area of the present study.

In contrast, for our eDNA study we sampled 22 sites along a non-overlapping 210 km stretch of the Teles Pires River in one year over the four water seasons, and we detected 10 orders, 40 families, 118 genera, and 265 species of fish. Of the 256 detected species, 108 were from the Teles Pires fishes in the Local reference library. Of the remaining 65 species of Teles Pires fishes in the Local reference library, we detected 28 sister species. This represents 74.6% of the species diversity detected by Ohara et al. [33] when using both the Midori2 and Local reference databases.

The 74.6% species detection level is the conservative lower limit, since not all species registered by Ohara et al. [33] for the lower reaches of the Teles Pires River were included in the Local reference database. It is also unclear if some of the species detected by Ohara et al. [33] do not occur in our sampled area due to regionalization of ichthyofauna within Amazonian River basins [58]. Furthermore, it is clear that no more than 50% of the ichthyofaunal diversity could be assigned to species level due to the deficiency of the reference database (Fig 3). Therefore, conservatively some 500 fish species are likely to occur in the area we surveyed using eDNA.

While 500 fish species may seem a large number, major Amazonian tributaries harbor anywhere from 502 to 1062 fish species [58]. The Tapajos River, which is formed by the confluence of the Juruena and Teles Pires rivers, has 522 registered species [58]. Additionally, a large number of species from the Amazon basin await description, and professionally conducted surveys almost invariably result in the discovery of new species, both previously described as well as new to science. For example, the multi-year ichthyological survey of the Madeira River prior to the construction of the Santo Antônio and Jirau hydroelectric dams resulted in a nearly 100% increase of species diversity—439 species listed in Torrente-Vilara [59] prior to the survey to more than 1000 species [58,60] following the survey.

The large number of detected species is a consequence of the primers used, and the availability of a comprehensive local reference database, whose development was possible due to the availability of genetic resources obtained through prior ichthyofaunal monitoring efforts using conventional sampling techniques. As expected, the use of a region-specific reference database significantly improves both detection efficiency and taxonomic resolution of eDNA-based studies. A customized reference library enhances the resolution of taxonomic assignments and increases overall detection efficiency [61]. Our results strongly support the effectiveness of eDNA metabarcoding for biodiversity monitoring in tropical freshwater systems, especially when paired with a well-curated local database.

Numerous studies have emphasized the scarcity of comprehensive 12S reference libraries, which results in few species-level identifications in eDNA studies [62,63]. These and our study highlight the urgent need for expanded efforts to enrich reference sequence databases, thereby improving the performance and reliability of eDNA-based surveys. One practical starting point for building a local reference library is to consult regional fauna checklists for the target area. In the absence of such lists, researchers can compile expected species inventories using resources like the Eschmeyer’s Catalog of Fishes [41] or from species distribution models such as in Map of Life (https://mol.org/). Subsequently, available sequences can be retrieved from GenBank or other platforms [64], or tissue samples from zoological collections can be obtained to generate new reference sequences [65].

Additionally, conducting traditional sampling in parallel with eDNA surveys provides valuable reference material and helps validate molecular detections [57]. Such integrative approaches are essential for the development of robust and taxonomically comprehensive databases that support accurate biodiversity assessments in megadiverse ecosystems like the Amazon.

## Conclusion

Environmental DNA studies are predicated on the specificity of primers, the taxonomic resolution of the amplified target region and reference databases. Studies focusing on DNA barcoding and early eDNA studies have proposed taxonomically informative target regions [25,26]. Subsequent studies investigated specificity of primers [18], recommending 12S and 16S primer sets. While specific, at least in the case of the standard 12S-V5 and MiFish primer sets, much of Neotropical fish diversity is not sampled since the primers do not incorporate priming site variation of the local ichthyofauna dominated by characiform fishes. The standard 12S-V5 and MiFish primer sets were developed for Paleoarctic fauna in general (12S-V5) and Paleoarctic and Asian fishes (MiFish). Numerous groups of Neotropical fishes have base pair mismatches at the 3’ end of these standard primers, and thus do not amplify, or do not amplify efficiently components of Neotropical ichthyofaunal. Consequently this generates false absences in local fish communities. A final key component of any eDNA study is the reference database. The Midori2 database includes a wide range of taxa from multiple vertebrate classes, making it a valuable tool when the objective is to assess broader vertebrate biodiversity. In our analysis, three vertebrate classes were detected using Midori2: Actinopterygii, Aves, and Mammalia (see Fig S1 and Table S3). Similar patterns were also reported by Ritter et al. [66], confirming that universal 12S primers—in our case the 12S-V5_mod primers, paired with broad databases can capture a wide range of vertebrate diversity. However, as emphasized earlier, the development and use of a Local reference database is critical for achieving accurate species-level identification—especially in highly diverse and understudied regions like the Amazon. When relying solely on Midori2, a high proportion of sequences are classified only at the Class level, due to the generic and incomplete nature of many entries (see Fig 3). In contrast, the combination of target-specific primers and a regionally curated database enables more precise taxonomic resolution, enhancing our ability to detect and monitor the rich Amazonian fish fauna with greater confidence and detail. This approach will be especially important for long-term monitoring and conservation of Amazonian aquatic biodiversity, which faces growing environmental pressures.

## Acknowledgments

We acknowledge Jaine Sousa and Gustavo Wolf from the Tropical Ichthyology Laboratory for their support with eDNA sample collection.

## Supporting information

**Table S1. List of species were identified to species level and deposited at the Instituto Nacional de Pesquisas da Amazônia (INPA) and Coleção Ictiológica do Acervo Biológico da Amazônia Meridional (ABAM-I), at Laboratório de Ictiologia Tropical (UFMT)**.(XLSX)

**Table S2 - Summary of Illumina reads and bioinformatic filtering.**

(XLSX)

**Table S3 - List of species by sampling point.**

(XLSX)

