## Supplementary figures and images for "Assessing fish diversity in the Amazon: The impact of primers and reference libraries in eDNA surveys"

### Suplemental Figure 1

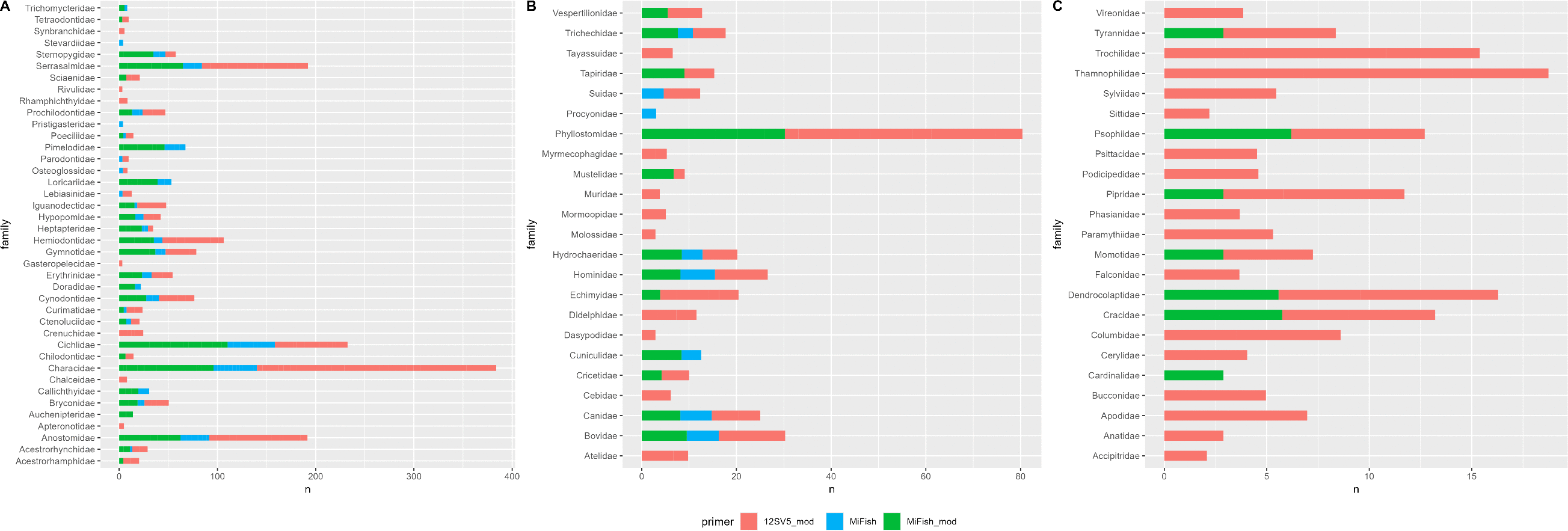
